# The immunomodulatory CEA cell adhesion molecule 6 (CEACAM6/CD66c) is a candidate receptor for the influenza A virus

**DOI:** 10.1101/104026

**Authors:** Shah Kamranur Rahman, Mairaj Ahmed Ansari, Pratibha Gaur, Imtiyaz Ahmad, Chandrani Chakravarty, Dileep Kumar Verma, Sanjay Chhibber, Naila Nehal, Shanmugaapriya Sellathanby, Dagmar Wirth, Gulam Waris, Sunil K. Lal

**Affiliations:** Virology Group, International Centre for Genetic Engineering & Biotechnology, New Delhi, India; Department of Microbiology and Immunology, H. M. Bligh Cancer Research Laboratories, Rosalind Franklin University of Medicine and Science, Chicago Medical School, North Chicago, Illinois, USA; Helmholtz Centre for Infection Research, Braunschweig, Germany; Department of Biomedical Science, Bharathidasan University, Trichy, India; Microbiology Department, Panjab University, Chandigarh, India; Career Institute of Medical & Dental Sciences and Hospital, Lucknow, India; School of Science, Monash University, Selangor DE, Malaysia

**Keywords:** Virus, Lipid raft, Flu, Influenza A Virus, Receptor, Neuraminidase, Hemagglutinin, CEA, CEACAM, CEACAM6, CD66c, IgG super family, Sialic acid

## Abstract

To establish a productive infection in host cells, viruses often use one or multiple host membrane glycoprotein as their receptors. For Influenza A virus (IAV) such a glycoprotein receptor has not been described, to date. Here we show that IAV is using the host membrane glycoprotein CD66c as a receptor for entry into human epithelial lung cells. Neuraminidase (NA), a viral spike protein binds to CD66c on the cell surface during IAV entry into the host cells. Lung cells overexpressing CD66c showed an increase in virus binding and subsequent entry into the cell. Upon comparison, CD66c demonstrated higher binding capacity than other membrane glycoproteins (EGFR and DC-SIGN) reported earlier to facilitate IAV entry into host cells. siRNA mediated knockdown of CD66c from lung cells inhibited virus binding on cell surface and entry into cells. Blocking CD66c by antibody on the cell surface resulted in decreased virus entry. We found CD66c is a specific glycoprotein receptor for influenza A virus that did not affect entry of non-IAV RNA virus (Hepatitis C virus). Finally, IAV pre-incubated with recombinant CD66c protein when administered intranasally in mice showed decreased cytopathic effects in mice lungs. This publication is the first to report CD66c (CEACAM6) as a glycoprotein receptor for Influenza A virus.

**Significance Statement:** Cells are enclosed by a semipermeable membrane that allows selective exchange of biomolecules between cells and their surroundings. A set of specialized proteins in this semipermeable membrane, work like gatekeepers to the cell and regulate entry of these biomolecules. One class of such surface proteins is termed as receptors. Viruses bind to one or more of these receptors and manipulate gatekeepers for their own successful entry into host-cells. A membrane protein that influenza A virus (Flu virus) uses for entry into the cells was not discovered till date. This study reports for the first time, a receptor for influenza A virus, that was sought after by researchers for decades. The viral receptor is a promising target that can be used to inhibit virus entry into host cells.

## Introduction

The outermost surface of mammalian cells typically bear a covering of branched sugar residues (oligosaccharides) that allow a wide range of interactions with different biomolecules (hormones, cytokines, growth factors etc.) of the matrix (1). These oligosaccharides are often linked to membrane proteins (glycoproteins) or lipid (gangliosides) at the cell surface. The type of sugar residues and their branching is responsible in deciding specificity of oligosaccharides towards biomolecules coming in contact with the cell surface. This interactions between oligosaccharides and biomolecules play a diverse physiological role and are important in maintaining communication and transport of molecules, between the cell and its surroundings (2). Many of these oligosaccharide chains bear sialic acid (SIA) at the termini, which serve as a regulators of molecular and cellular interactions (3).

In general, viruses often breach this communication and bind to terminal SIA as the first step to invade cells (4). This less specific interaction of viruses with terminal SIA is followed by a more specific interaction of viral spike proteins with a subset of host glycoprotein receptors that effectively accompany the virus into cells and drive the ingested cargo to destined endocytic pathways or intracellular routings (5).

The mechanism of Influenza A Virus (IAV) entry still remains elusive. Information on a host glycoprotein receptor that can pull virus into the cell is largely unknown. Most of the published literature on influenza entry are centric to the early attachment factor viz. oligosaccharides. The importance of the role of SIA in IAV entry has been documented as early as 1959 (6) however, there are reports showing IAV entry into host cells even in the absence of SIA (7, 8, 9). Interestingly, De Vries *et. al*. and Chu *et. al*. in their studies suggested that while SIAs (α2–6, α2–3) may help in the attachment of virus, a specific subset of glycoprotein receptor is necessary for virus entry which is yet unidentified (8, 9). Over the years, a few membrane glycoproteins like EGFR (epidermal growth factor receptor), L-SIGN (liver/lymph node-specific intracellular adhesion molecules-3 grabbing non-integrin) and DC-SIGN (Dendritic Cell-Specific Intercellular adhesion molecule-3-Grabbing Non-integrin) were reported to facilitate viral attachment and entry but were not designated as a receptor for IAV possibly due to lack of evidence that they play a major role in IAV uptake (7, 10). Moreover, the evidence of their physical interaction with IAV spike proteins, Hemagglutinin (HA) and Neuraminidase (NA), at the cell surface or inside the cell was not well characterised. Generally, the role of glycoproteins EGFR, L-SIGN and DC-SIGN in the cellular uptake of virus is believed to be a low-specificity phenomenon since these surface proteins facilitate uptake of a number of viruses (11). Also, a comparative study showing specificity of these glycoproteins towards IAV and not other viruses was not demonstrated.

In this report, we have identified a glycoprotein receptor for IAV entry into lung epithelial cells. Earlier, we published a detailed account of an interaction between NA and a host membrane glycoprotein CD66c, validated for a variety of different IAV isolates (12). We now report CD66c as the first glycoprotein receptor candidate for IAV entry into host cells. CD66c aka Carcinoembryonic Cell Adhesion Molecule 6 (CEACAM6) is a GPI-anchored, raft associated, highly sialylated membrane glycoprotein of the immunoglobulin superfamily (IgSF).

To systematically validate CD66c as a receptor, we overexpressed this molecule on the surface of human lung cells and studied IAV binding and entry. Cells overexpressing CD66c on cell surface showed an increase in virus binding on the cell surface and subsequently increased entry into cells. Besides lung cell line (A549), the effect of CD66c on virus entry was further tested in mouse fibroblasts cells (NIH3T3), chinese hamster ovary cells (Lec2 CHO) and human embryonic kidney cells (HEK293). On the contrary, when CD66c expression levels were reduced by siRNA-mediated knockdown, we observed a significant decrease in viral binding and entry into lung cells. To further investigate the role of interaction, between viral NA and host CD66c at the cell-surface, in virus entry, we performed antibody-mediated receptor-blockade experiment. For this experiment, surface receptor CD66c was masked by incubating a monolayer of A549 lung cells with anti-CD66c monoclonal antibody (mAb) prior to virus binding to this monolayer. Masking of receptor CD66c by antibody inhibited virus entry probably due to a restricted interaction between the viral spike NA and putative receptor CD66c. Having validated CD66c as a receptor for IAV, it was important to compare the binding capacity of CD66c, towards virus binding and entry, with respect to other host glycoproteins (EGFR, DC-SIGN) that had been reported to facilitate virus binding and entry (7, 10). We noticed, a significant increase in virus binding and entry into lung cells overexpressing CD66c however cells overexpressing EGFR, DC-SIGN showed a modest increase in virus binding and entry, under similar experimental conditions. We also found that overexpression of CD66c did not affect the expression levels of EGFR, DC-SIGN and vice versa. Additionally, we carried out an important experiment to validate CD66c as a specific receptor for the influenza virus. While glycoprotein CD66c was found necessary for the uptake of influenza virus by cells, it did not affect the entry of another non-IAV RNA virus (Hepatitis C virus). Finally, to confirm our hypothesis, we pre-incubated IAV particles with biologically active human recombinant CD66c protein (expressed in mouse myeloma cell line) prior to infecting BALB/c mice. Interestingly, we observed a significant decrease in infection and cytopathic effects in mice lungs presumably due to masking of the NA spike protein with rCD66c protein.

This study on identification of CD66c as a receptor for the influenza A virus has 124 potential to further our knowledge on the mechanism of virus entry. Additionally, this newfound receptor brings to attention a new target for anti-viral interventions. Since CD66c is also an active immunomodulatory molecule (present on T cells, B cells and neutrophils) this finding may lead to exploration of early immunomodulatory events associated with the interaction between viral NA and CD66c.

## Results

### Overexpression of CD66c on human lung cell surface resulted in increased virus binding

The physical interaction between influenza NA and CD66c of virus-infected cells was confirmed earlier by co-immunoprecipitation assays (12). Here, we performed experiments to quantify virus binding on the surface of cells overexpressing CD66c glycoproteins. A549 lung cells transiently transfected with untagged CD66c gene showed a rise in the expression of this molecule at the host cell surface when monitored by flow cytometry (**Figure 1a**). Further, lung cells overexpressing CD66c showed significantly increased virus binding on cell surface. Increase in virus binding was detected by quantitatively probing NA, on the host cell surface. Our results showed an increase in viral NA on the surface of CD66c overexpressing cells, indicating higher virus binding (**Figure 1b**). Flow cytometry results showed that among cells with endogenous level of CD66c, ~18% of the total cell population had virus bound to cell surface. In contrast, CD66c overexpressing cells showed that approximately 80% of total cell population had virus bound to the cell surface (**Figure 1b**). We further carried out experiments to investigate whether the NA present on virus envelope colocalized with the putative receptor CD66c at the cell surface. For this experiment, we let IAV bind over a monolayer of human lung cells and subsequently stained cells with a mixture of two antibodies, anti-NA and anti-CD66c. Confocal microscopy revealed colocalization of CD66c with viral NA at the cell surface (**Figure 1c**). The CD66c (red) and viral NA (green) bound to the cell surface were clearly visible on merged view (yellow) (**Figure 1c**).

**Figure 1:**
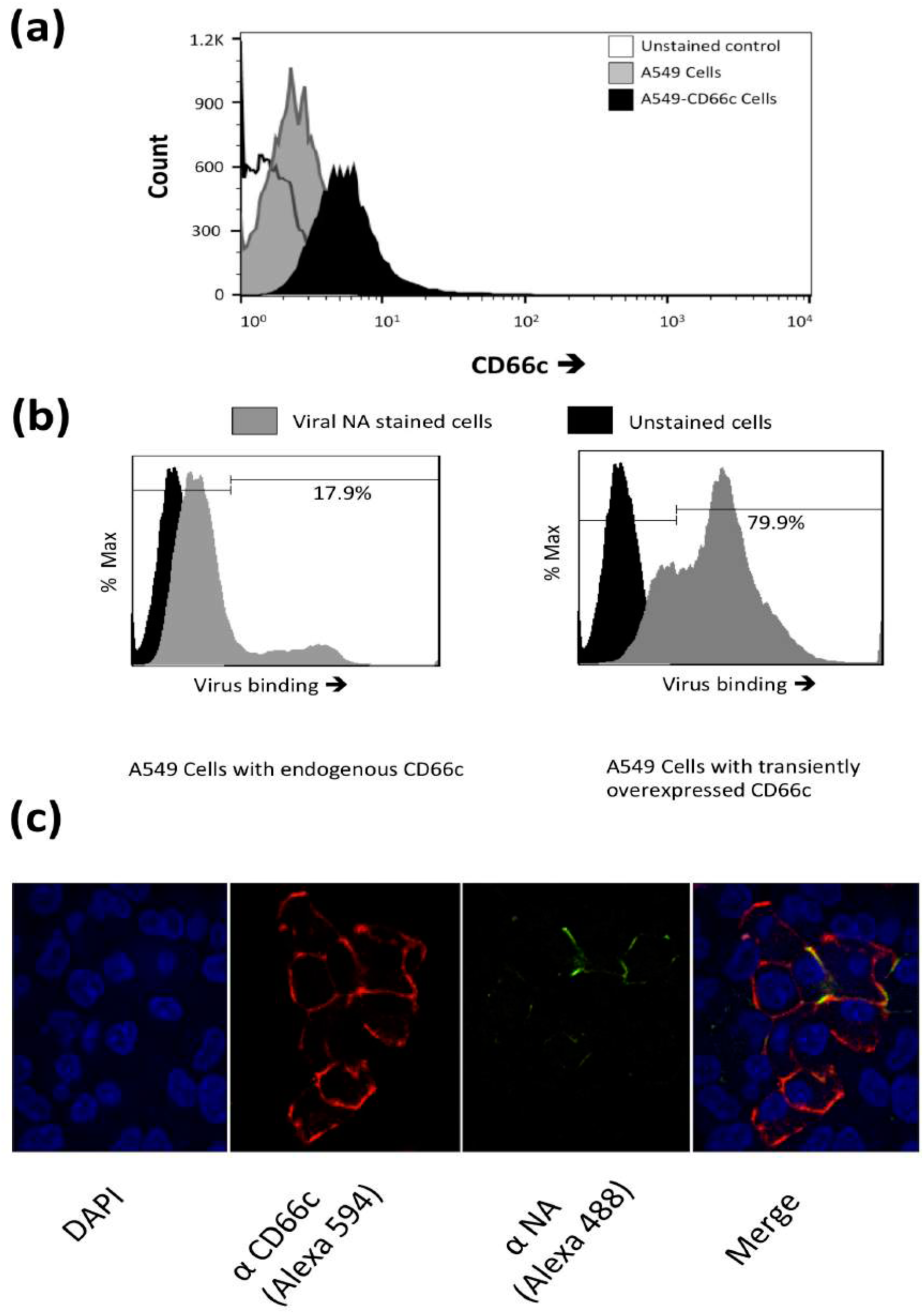
**(a): Flow cytometry analysis of surface expression of CD66c in A549 cells.** Cell surface expression of untagged CD66c was determined using flow cytometer, where cells were stained for CD66c (Alexa-594). Cell population transiently overexpressing CD66c shows signal for higher expression of CD66c on the cell surface (black) as compared to A549 cells (grey). The unstained control cells are shown in white. **Figure 1 (b): Flow cytometry analysis of virus binding on A549 cells and CD66c overexpressing A549 cells**. 5 multiplicity of infection (m.o.i.) of PR8 virus binding to a monolayer of A549 cells was measured by determining signals corresponding to stained NA protein (Alexa-488) of the virus on host cell surface. In figure, black curve denotes signal of unstained cells while grey curve depicts the signal for viral NA corresponding to the virus binding on the cell surface. The left panel shows that virus is binding to ~ 18% of A549 cell population. However, the right panel shows that PR8 virus binding is increased to more than 80% of CD66c overexpressing A549 cell population. **Figure 1 (c): Demonstration of virus binding to A549 cell surface, through immunofluorescence assay and confocal microscope**. To a monolayer of A549 cells, binding of 5 multiplicity of infection (moi) of PR8 virus was observed under confocal microscope (A1R; Nikon, Tokyo, Japan) at 60X magnification, after extracellular immune-fluorescence-assay (IFA) staining of viral NA and host CD66c protein. Panels can be numbered 1 to 4 from left to right. Panel 1 shows cell nuclei stained with DAPI in cells of the selected microscopic field. Panel 2 shows that, CD66c (red) being a membrane protein is mainly at the periphery of cells. Panel 3 shows that virus is present at the surface of A549 cells (green), as cells were fixed just after their brief binding on cell. In Panel 4 we see the colocalisation (yellow) of NA protein of PR8 virus (green) with host membrane protein CD66c (red) at the periphery of cells.

### Cells overexpressing CD66c showed increased virus entry

We further sought to determine if CD66c overexpression in host cells increased virus entry. We quantified mRNA levels of viral NP and M1 after harvesting PR8 infected A549 lung cells, approximately 8 hours post-infection, h.p.i (one life cycle). Cells overexpressing CD66c showed 6-8 fold increase in the levels of M1 and NP mRNA (**Figure 2a, b**). We also measured viral NP protein in CD66c expressing A549 cells that showed 7 fold increase in its expression levels (**Figure 2c-f**). To demonstrate that the NA-CD66c interaction affected virus entry in other types of cells as well, we performed similar experiments using the mouse embryonic fibroblast cell line NIH3T3. CD66c expressing NIH3T3 cells (NIH3T3-CD66c cells) showed an increase in NP mRNA levels as against the NIH3T3 cells (**Figure 3a**). Likewise we checked the virus binding and entry in CD66c overexpressing Lec2CHO cell lines. Lec2CHO cell lines transiently overexpressing CD66c favored virus binding on cell surface and subsequent uptake of the virus into cells (**Figure 3b, c**). HEK cells overexpressing CD66c cells showed an increase in virus entry, which was monitored by increase in the NP mRNA levels (**Figure 3d**).

**Figure 2:**
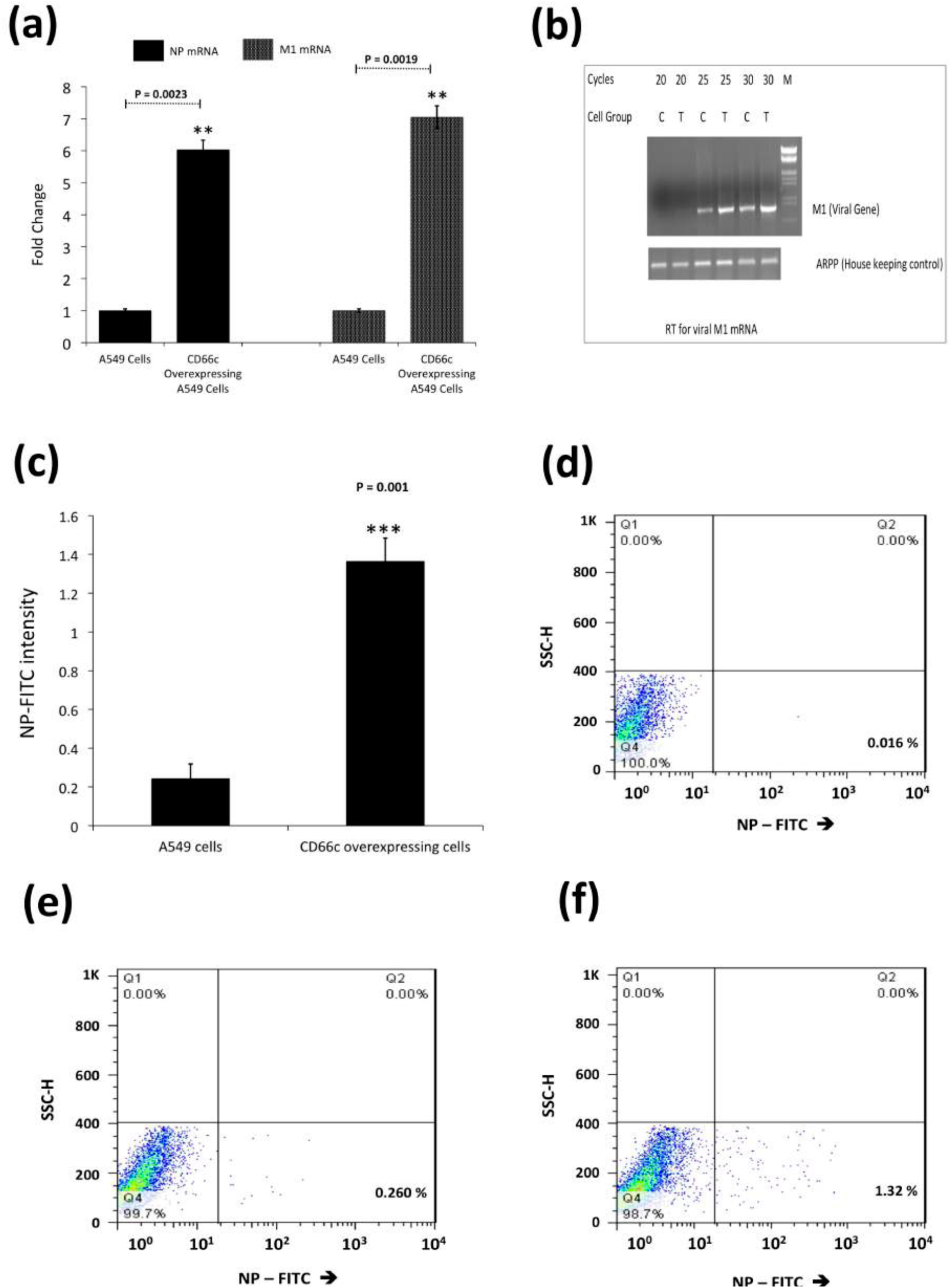
**(a): Real time quantification of NP and M1 mRNA in viral infected A549 cells.** The untagged CD66c expressing plasmids were transfected into A549 lung cell lines. 48 hour post transfection, cells were infected with 1.0 multiplicity of infection (m.o.i.) A/PR8/34 influenza virus. 8 hours post infection (8 h.p.i) infected cells were harvested for mRNA isolation and subsequent quantification. Levels of two viral mRNA NP and M1 were quantified to compare the viral load in infected lung cells. Figure shows higher mRNA levels of NP (left two bars) and M1 (right two bars) in cells overexpressing CD66c. Each bar represents mean of five independent experimental readings. **(b): Semi quantification of viral M1 by RT-PCR**. For the above mentioned reaction condition (8 h.p.i, and 1 m.o.i.) viral M1 mRNA was also measured semi-quantitatively. A549 cells are denoted with ‘C’ before respective lanes and likewise A549 cells having overexpressed CD66c with ‘T’. Cycles are the number of PCR cycles. **(c): Flow cytometric analysis for virus load in A549 infected cell**: Following similar gene expression and viral infection conditions (8 h.p.i,), in another set of experiment, A549 cells were harvested to quantify protein level of viral NP through flow cytometry, in cells infected with 0.5 m.o.i. of A/PR8/34 influenza virus. NP expression levels in cells were measured through intracellular staining by NP-FITC conjugated antibody. The bar diagram shows rise in viral NP in CD66c overexpressing A549 cells (right) than that in A549 cells (left). Each bar represents mean of three independent experimental readings. Data represent mean values of at least three independent experiments ± SD. Statistical significance was assessed by student’s t-test, (*) for p ≤ 0.05, (**) for p ≤ 0.01 and (***) for P ≤ 0.001. **(d):** Representative FACS snapshot of unstained A549 cells showing no signal corresponding to NP-FITC in lower-right quadrant. **(e):** A549 cells show some signal corresponding to NP-FITC in lower-right quadrant. **(f):** A549 cells overexpressing CD66c is shown to have increased NP-FITC signal in lower-right quadrant.

**Figure 3:**
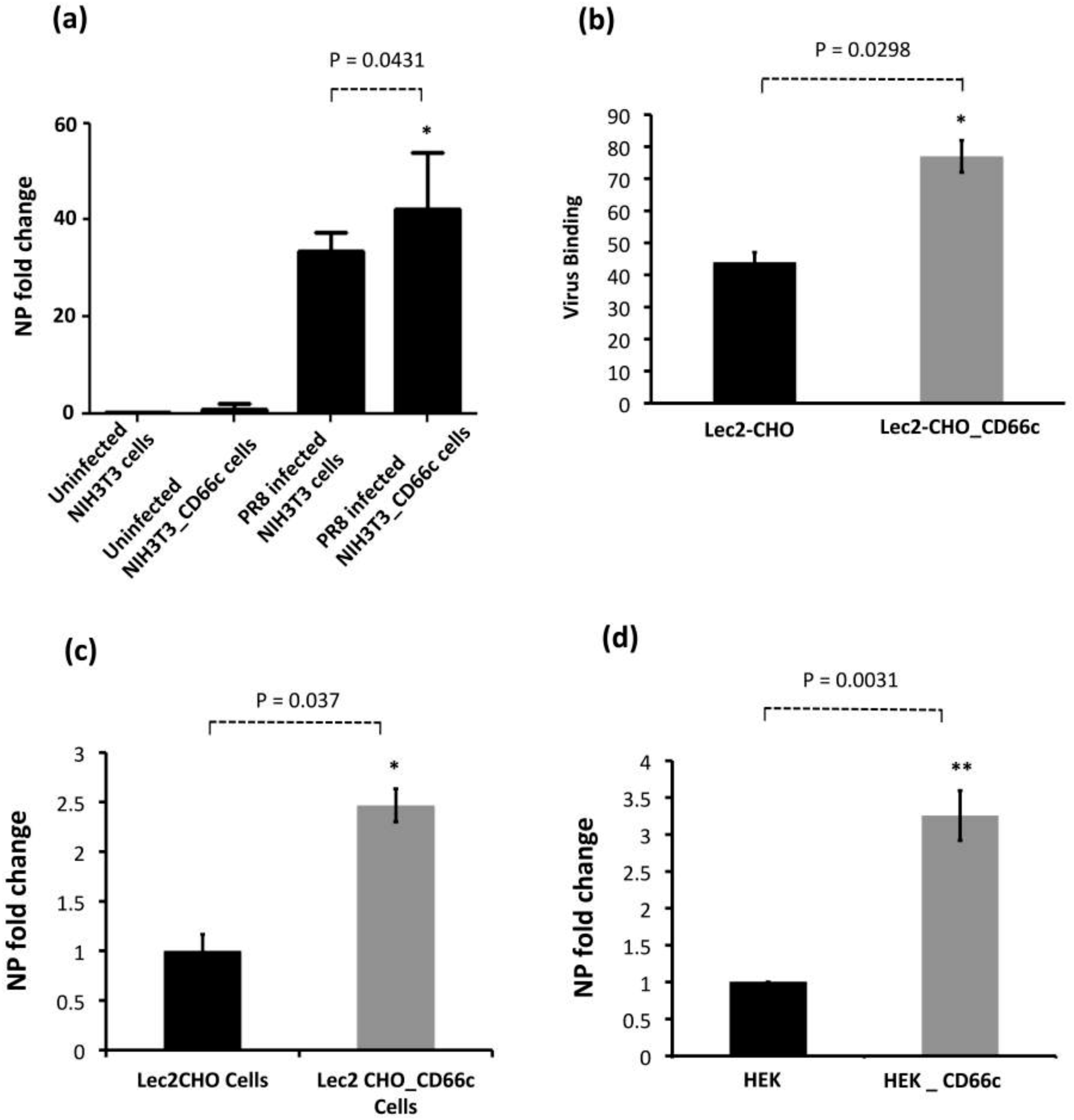
**Demonstration of rise in viral uptake in CD66c overexpressing NIH 3T3 and Lec2-CHO cell lines. (a):** The level of infection as probed by the NP mRNA fold-change is shown as bars. The bar in the right end shows increased influenza infection in CD66c transfected NIH3T3 cell lines. **(b):** Lec2 CHO-CD66c is CD66c overexpressing Lec2 CHO cell lines. The right bar shows greater virus binding on the surface of cells overexpressing CD66c. **(c):** The right bar in the figure shows increased level of viral NP mRNA Lec2 CHO-CD66c cells (right bar) as compared to Lec2 CHO cells (left bar) suggesting increased virus entry. **(d):** The left bar shows levels of mRNA corresponding to lower infection level in HEK cells and the right bar shows an increase in infection in CD66c overexpressing HEK cells.

### siRNA knockdown of CD66c inhibited virus binding on cell surface and subsequent entry into lung cells

After having conducted overexpression studies with CD66c, we carried out siRNA experiments to silence the expression of this molecule and subsequently studied virus binding and entry into lung epithelial cells. For virus binding experiments cells incubated with virus for a brief period of 5 minutes were stained extracellularly for receptor CD66c (green) and Neuraminidase (red) and observed under fluorescent microscope. We found that siRNA mediated knockdown of CD66c expression in A549 lung cells resulted in the inhibition of virus binding on the cell surface (**Figure 4a**). Cells treated with CD66c siRNA showed poor expression of CD66c as shown by a weak green signal in uninfected cells (UI) and 5 minutes post-infected cells (5’), as against control siRNA treated cells (**Figure 4a**). Also, virus binding was not observed on the surface of cells silenced for CD66c, as observed by a reduced red signal for neuraminidase (Lower two panels, **Figure 4a**). In contrast, mock siRNA treated cells showed significant endogenous expression levels of CD66c (green) both in viral infected (5’) and uninfected cells (UI). As expected, these cells upon infection showed significant viral binding on the cell surface, which was probed by viral NA (red). Accordingly, these cells showed significant colocalization of NA (red) with CD66c (green) in merged fields (yellow) (**Figure 4a**). Altogether, we observed that cells with endogenous level of CD66c showed significant virus binding at the cell surface however cells silenced for CD66c did not show any visible virus binding on the cell surface due to absence of surface receptor CD66c (**Figure 4a**). To validate our immunofluorescence assay (IFA) data we performed a western blot analysis to study the effect of siRNA-mediated-silencing of CD66c in virus entry (**Figure 4b**). The siRNA treated lung cells that showed complete loss of CD66c expression, consequently demonstrated inhibition of viral infection as was evident from poor expression of the viral protein NP (**Figure 4b**). Additionally, using western blot analysis, effect of CD66c silencing on expression levels of membrane protein EGFR and DC-SIGN were evaluated. We found that siRNA mediated silencing of CD66c did not suppress the expression of EGFR and DC-SIGN (**Figure 4b**). More importantly this data also suggested that expression of CD66c in A549 lung cells did not have any effect on the expression levels of EGFR and DC-SIGN.

**Figure 4 (a):**
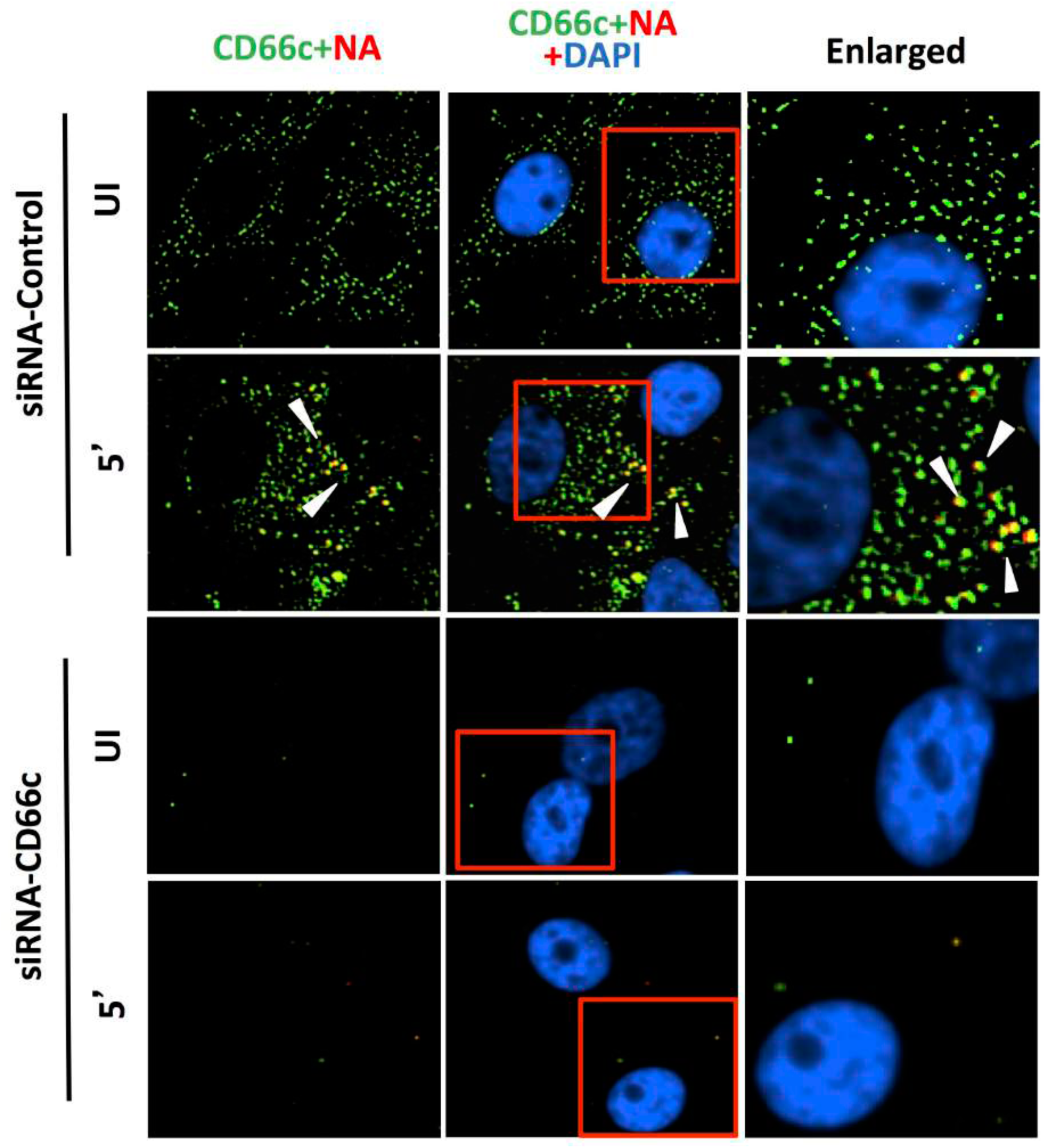
**Knockdown of CD66c shows inhibition of virus binding on cell surface under fluorescent microscope.** Here, UI denotes uninfected cells; 5’, Cells after 5 minutes of virus binding to them. Figure shows that A549 cells treated with siRNA control (negative) do not inhibit expression level of CD66c (green) and therefore binding of 5 MOI of PR8 viruses on cell surface can be seen. Figure shows colocalisation between NA (red) and CD66c (green) in merged view (yellow) (second panel from top, pointed with white arrow). Amount of colocalization between NA and CD66c and absence of NA (red color) in control siRNA treated cells signifies possible colocalisation between CD66c and NA at the host cell surface. The lowest two panels show diminished green signals in cells treated with CD66c siRNA suggesting a poor expression of CD66c protein (green). Consequently, virus binding on siRNA-CD66c treated cells is not seen, as evident by the absence of any green (CD66c) or red (NA) signal (bottom panel).

**Figure 4 (b):**
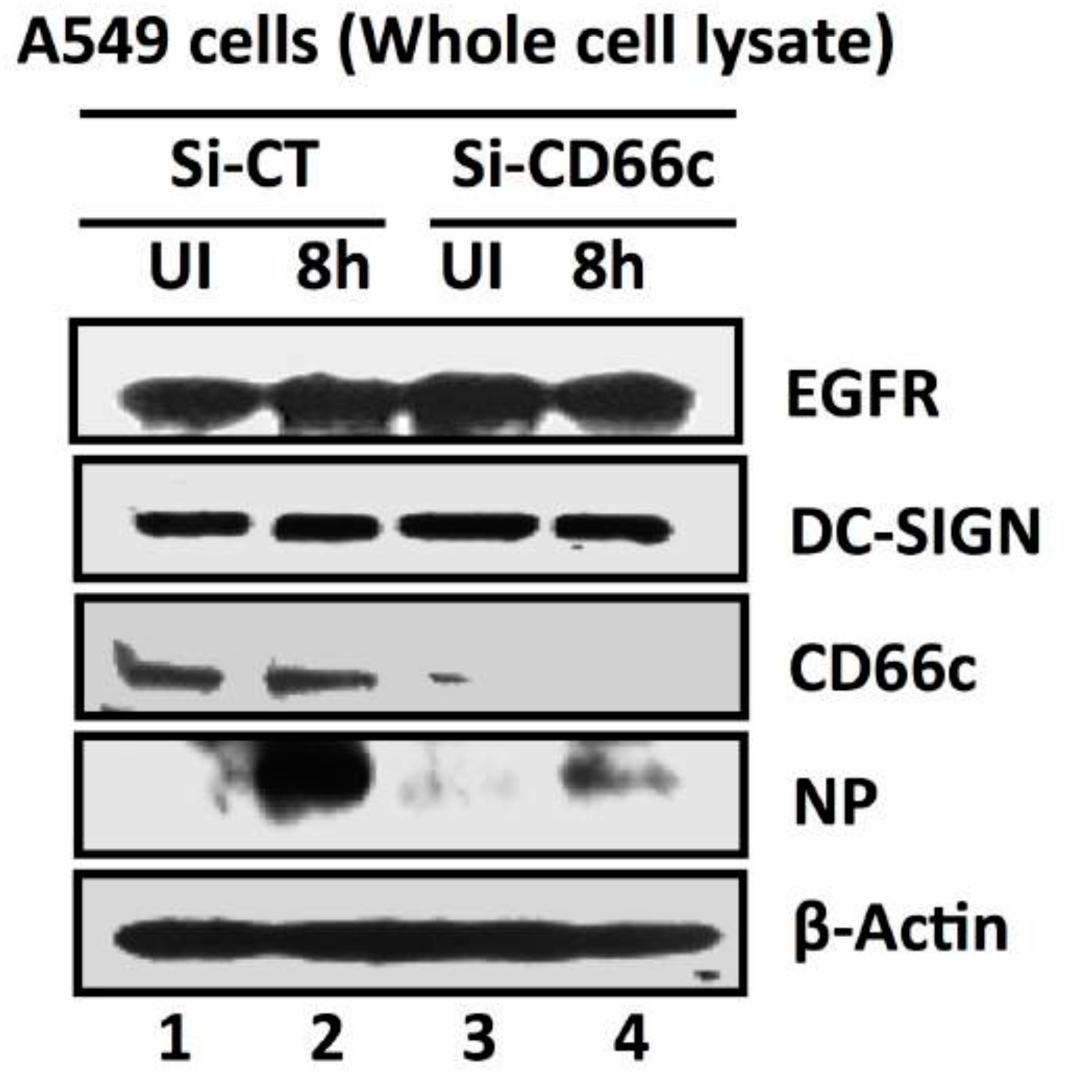
**siRNA-mediated knockdown of CD66c shows inhibition of virus entry into lung-cells infected with 1 m.o.i. of PR8 virus, through western blot analysis.** Here, Si-CT denotes control siRNA treated cells; Si-CD66c is cells treated with CD66c siRNA. 8h is cells harvested after 8 hours of infection (one life cycle of IAV), UI, is uninfected cells. A549 cells treated with siRNA shows complete knock down of the receptor molecule CD66c (third panel from top) while control siRNA treated cells do not show any reduction in expression of CD66c (left two wells of third panel). The fourth panel from top shows level of A/PR8/34 influenza virus entry in cells determined by expression levels of viral NP protein. The second well from left shows significant expression level of viral NP protein in cells treated with control siRNA, eight hours after infection. In contrast, the right most well shows a marked reduction of IAV entry in cells treated with CD66c siRNA, as determined by low expression level of viral protein NP eight hours after infection. Conclusively, virus entry was inhibited in absence of CD66c (CD66c siRNA treated cells). CD66c siRNA treated cells do not show any noticeable change in expression levels of EGRF (top most panel) and DC-SIGN (second panel from top). For loading control β-Actin was probed (the bottom panel).

### Antibody-mediated masking of receptor CD66c at cell surface inhibited virus entry

From our previous publication, it was established that NA interacts with CD66c inside the host cell and overexpression of CD66c influenced cell survival pathways (PI3K-Akt), with subsequent increase in viral load in infected cells (12). In that context, here a decrease in viral load in cells knocked down for CD66c (**Figure 4a**) may be implicated due to a corresponding down modulation of cell survival pathway (PI3K-Akt). Therefore this result (**Figure 4a**) is not sufficient to claim CD66c is a receptor. To validate CD66c is a receptor for viral entry, we need evidence to demonstrate effect of direct interaction, between CD66c and NA at the host cell surface, in virus entry. Therefore, we conducted the receptor-blockade experiment. An experiment demonstrating that a disruption of physical interaction between host CD66c and viral NA at the cell surface could affect viral entry. Initially, we performed antibody-mediated receptor blockade experiments on CD66c overexpressing NIH3T3 (NIH3T3-CD66c) cells. For this experiment, a monolayer of NIH3T3-CD66c cells when treated with mAb anti-CD66c at increasing concentrations of 1.0μg/mL, 1.5μg/mL, 2.0μg/mL and 8.0μg/mL, preceding virus infection. This experiment revealed significant decrease in virus infection in a dose dependent manner. The virus entry levels in these cells were determined by the mRNA levels of viral NP (**Figure 5a**). Likewise inhibition of virus entry was observed by determining expression levels of viral NP protein in NIH3T3-CD66c cells, when treated with increasing concentrations of anti-CD66c mAb prior to infection (**Figure 5b**). From these results, lower levels of NP (mRNA and protein) in anti-CD66c mAb treated cells suggested that masking of CD66c on host cell surface by mAb reduced its access to NA spike protein present on the envelope of the infecting virus particles thus inhibiting virus binding and uptake. For conclusive validation of our hypothesis, we finally performed mAb mediated receptor blockade experiment in A549 lung cells with endogenous (not overexpressed) levels of CD66c on the cell surface. In a control experiment we also tested the effect of mock antibody (IgG isotype) binding on the A549 cell monolayer as against CD66c binding (**Figure 6a, b**). We observed a corresponding decrease in virus entry when cells were treated with mAb anti-CD66c at the respective concentrations of 1.0 μg/mL, 1.5μg/mL, 2.0μg/mL, 4.0μg/mL and 8.0 μg/mL (**Figure 6c**). Inhibition of virus entry was demonstrated by measuring a corresponding reduction in expression levels of viral NP protein in infected cells by flow cytometry (**Figure 6d-g**). The dose dependence of mAb anti-CD66c in the inhibition of viral entry was also confirmed by quantitating another viral protein, M1 in the corresponding cells (**Figure 6h, i**). One set of the antibody mediated receptor blockade experiment was also studied under confocal microscopy, which showed similar inhibition of virus entry into A549 cells that were treated with 4.0 μg/mL of anti-CD66c mAb (**Figure 6j**).

**Figure 5:**
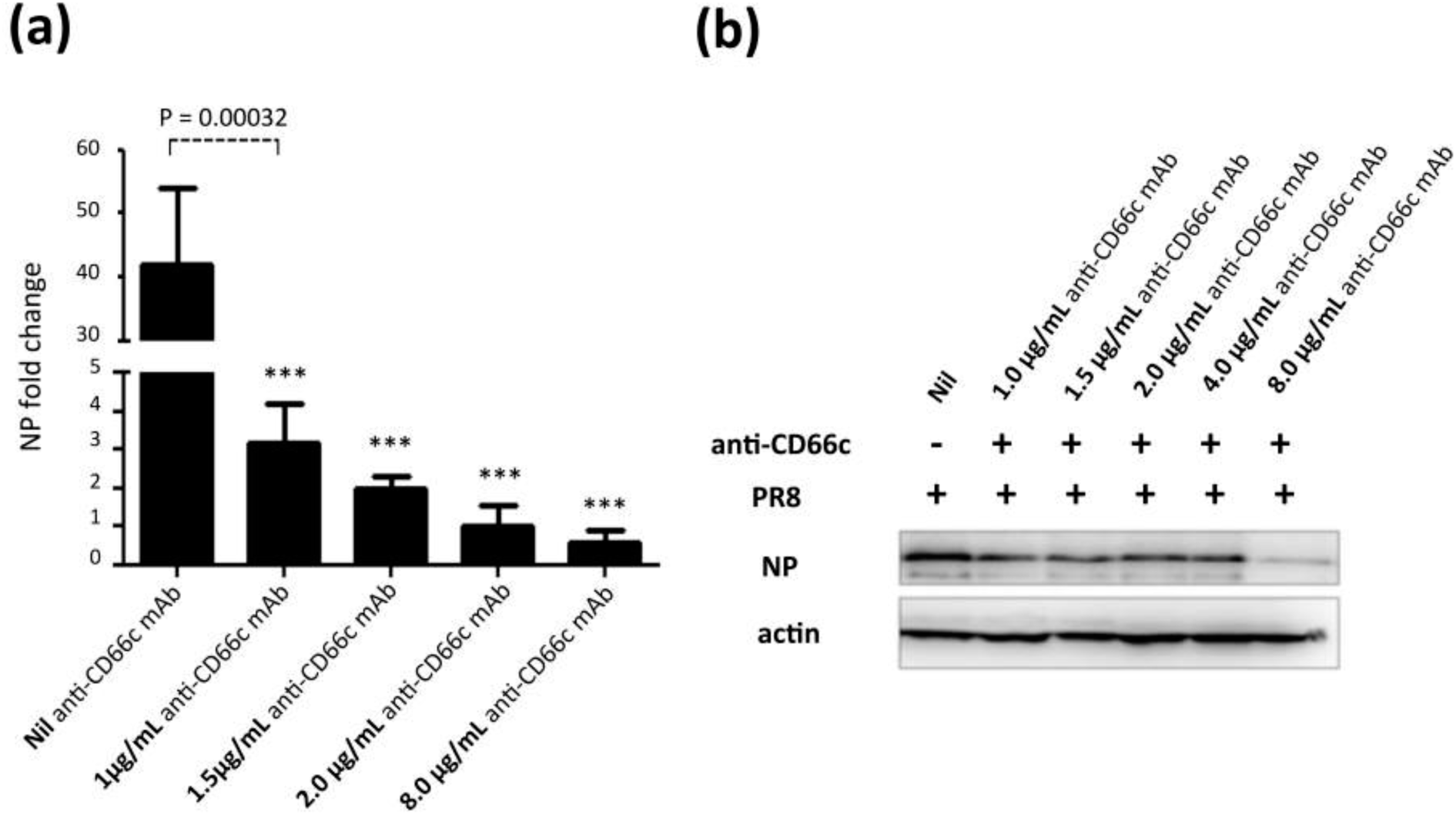
**Antibody mediated receptor blockade experiments in CD66c overexpressing cells. (a):** The figure shows level of A/PR8/34 influenza virus entry in a monolayer of NIH3T3-CD66c cells when treated with mAb anti-CD66c prior to infection. From left to right, the first bar in the figure represents levels of viral NP mRNA (a measure of virus entry) in untreated NIH3T3-CD66c. Bars second to fifth from left show levels of viral NP mRNA in virus-infected cells treated with mAb anti-CD66c at the concentration of 1.0 μg/mL, 1.5 μg/mL, 2.0 μg/mL and 8.0 μg/mL respectively. Conclusively, the data show decrease in virus entry in cells treated with anti-CD66c in a dose dependent manner. Data represent mean values of at least three independent experiments ± SD. Statistical significance was assessed by student’s t-test (GraphPad), (*) for p ≤ 0.05, (**) for p ≤ 0.01 and (***) for P ≤ 0.001. **(b):** A western blot showing expression levels of viral NP protein in virus-infected NIH3T3-CD66c when treated with corresponding concentrations of mAb anti-CD66c 1.0 μg/mL, 1.5 μg/mL, 2.0 μg/mL, 4.0 μg/mL and 8.0 μg/mL, prior to viral infection. The inhibition of virus entry in anti-CD66c mAb treated cells was significant at a concentration of 8.0 μg/mL of CD66c.

**Figure 6:**
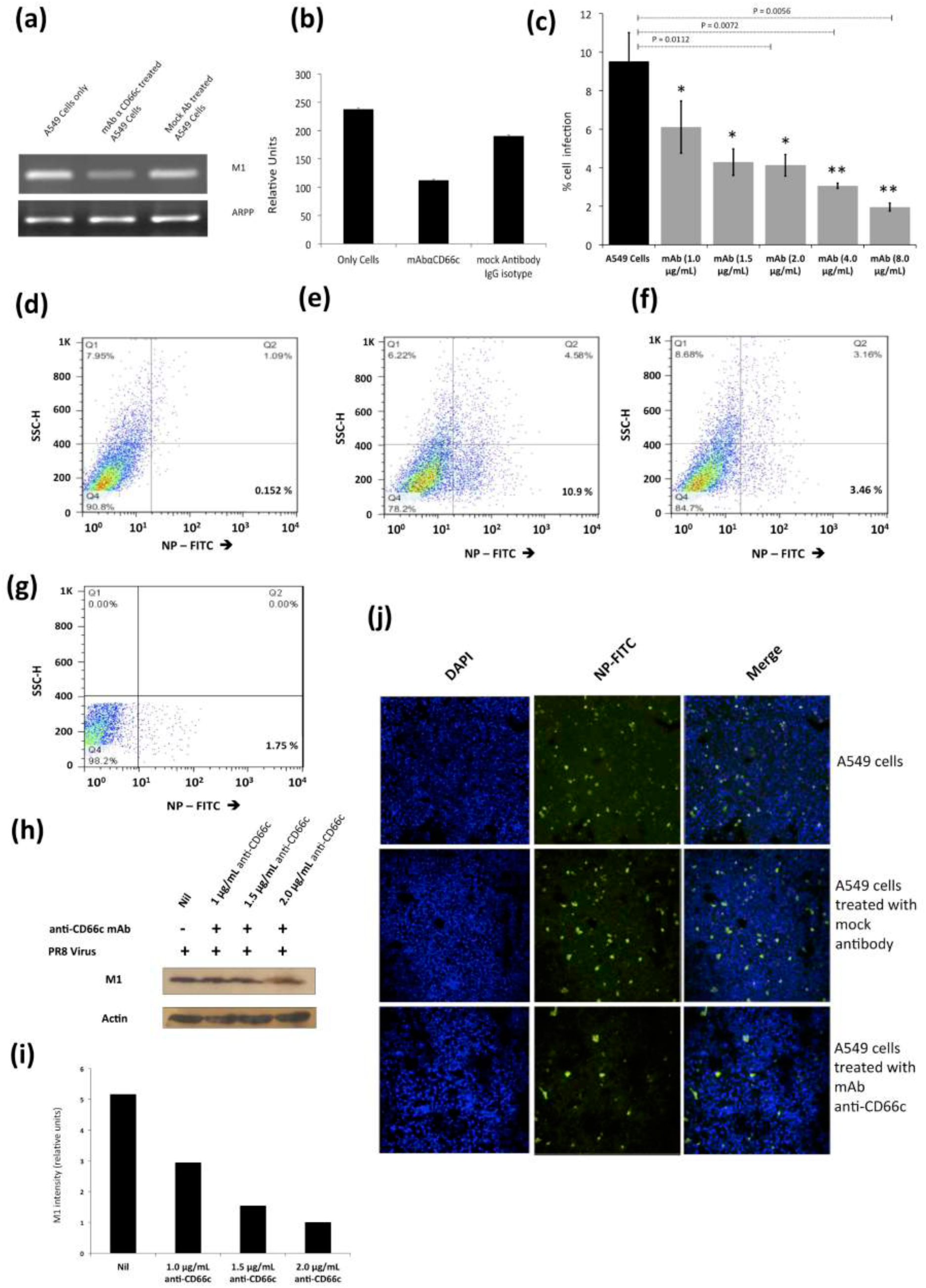
**Antibody mediated receptor blockade experiments in A549 cells expressing endogenous levels of CD66c.** Figure shows A/PR8/34 influenza virus infection levels in lung A549 cells (24 h.p.i.), when incubated with increasing concentrations of anti-CD66c mAb prior to infection. **(a):** Semi quantification of viral M1 mRNA shows significant difference in virus entry. Left well shows M1 mRNA level in A549 cells, the middle well shows that in cells incubated with mAb anti-CD66c. And the right well shows M1 mRNA level in cells incubated with mock antibody (IgG isotype antibody) prior to viral infection. **(b):** Densitometry analysis of image (a). The cells treated with anti-CD66c showed reduced virus entry as against the untreated and IgG isotype antibody treated cells. The expression level of housekeeping gene acidic ribosomal phosphoprotein (ARPP) is not affected in any of these cells. **(c):** The black bar represents A549 cell populations expressing viral NP protein (a measure of virus entry) after infection without any anti-CD66c treatment. Whereas grey bars (from left to right) represent the same in A549 cells treated with anti-CD66c mAb, prior to infection, at the concentrations of 1.0 μg/mL, 1.5 μg/mL, 2 μg/mL, 4 μg/mL and 8 μg/mL respectively. Here, the population of virus-infected cells expressing viral NP protein determines extent of virus entry. Conclusively, the bar diagram shows that cells treated with anti-CD66c mAb showed inhibited virus entry. The inhibition of the virus entry varied in a dose (of anti-CD66c) dependent manner. Each bar represents mean of three independent experimental readings. Data show the mean percent infection (± SD) from 4 independent experiments. (*, P < 0.05; **, P < 0.01) Figures (**d-g**) are representative snapshots of NP stained cells from flow cytometry. **(d):** Unstained A549 cells showing baseline signal in lower-right quadrant, in 0.0152 % cell population. **(e):** A549 cells showing good signal corresponding to NP-FITC in lower-right quadrant, in 10.9 % cell population. **(f):** shows A549 cells incubated with 4.0 μg/mL of mAb anti-CD66c with relatively reduced NP-FITC signal in the lower-right quadrant, in 3.46 % cell population. **(g):** A549 cells incubated with 8.0 μg/mL of mAb anti-CD66c is showing reduced NP-FITC signal in lower-right quadrant, in 1.75 % cell population. **(h): Western blot analysis of infection in A549 cells incubated with anti-CD66c mAb.** The figure shows levels of viral M1 protein expression in virus-infected cells treated with anti-CD66c mAb at a concentration of 1.0 μg/mL, 1.5 μg/mL and 2.0 μg/mL, prior to infection by the virus. It shows a corresponding decrease in the level of viral M1 protein expression 24 h.p.i in a dose (of anti-CD66c) dependent manner, suggesting inhibition of virus entry in anti-CD66c treated cells. **(i):** Densitometry analysis of the western blot shown in (h). From left to right are the bars representing the level of M1 expression in untreated cells and in those treated with mAb anti-CD66c at a concentrations of 1.0 μg /mL, 1.5 μg/mL and 2.0 μg/mL respectively. **(j): Immunofluorescent assay (IFA) and confocal microscopic analysis of virus entry in A549.** The figure has three panels from top to bottom each with three images. From left to right in each panel, we have A549 A/PR8/34 virus-infected cells showing nuclei stained with DAPI (*blue*), viral NP stained with FITC conjugated primary antibody (*green*) and a superimposed image of the first two figures showing both channels (blue and green). Here, levels of NP in A549 cells 24 h.p.i are a measure of virus entry in cells. Uppermost panel shows infection level in A549 cells. The middle panel shows infection level in cells that are incubated with mock antibody (IgG Isotype control) prior to virus infection on the monolayer. The bottom panel shows reduced infection in cells that were incubated with 4 μg/mL anti-CD66c mAb before viral infection.

### Virus binding and entry in cells overexpressing CD66c, EGFR and DC-SIGN

A few reports earlier showed that overexpression of two membrane proteins (EGFR and DC-SIGN) resulted an increase in virus binding and entry into mammalian cells. Therefore we conducted an experiment to compare virus binding and entry in cells overexpressing CD66c, EGFR and DC-SIGN respectively. For this experiment we checked the extent of virus binding on the lung cells, at the endogenous and overexpressed levels of these host membrane glycoproteins (EGFR, DC-SIGN and CD66c). We reasoned that a genuine receptor upon overexpression in lung cells should exhibit significant increase in virus binding on the cell surface, whereas a weak receptor candidate upon overexpression should display a modest increase in virus binding. After allowing IAV virus to bind to cell monolayers, we examined the cultures under a fluorescence microscope to monitor the membrane glycoproteins (green) and viral NA (red). We did not notice significant increase in virus binding on lung cells overexpressing EGFR as against cells with endogenous levels of EGFR (**Figure 7a**). Also, the viral spike protein NA (red) did not show possible colocalization with EGFR. Similarly, there was no significant increase in virus binding on lung cells overexpressing DC-SIGN as compared to cells with endogenous levels of DC-SIGN (**Figure 7b**). However, when we analyzed cells with endogenous level of receptor CD66c it showed significant virus binding (**Figure 7c**), the yellow signal in these cells suggested strong colocalization of NA (red) with receptor CD66c (green). Additionally, upon CD66c overexpression lung cells showed a further increase in virus binding on cells as against cells with endogenous CD66c level (**Figure 7c**) Altogether, the corresponding increase in yellow spots in cells overexpressing CD66c signifies the higher binding capacity of CD66c (green) towards Influenza A virus (NA) at the cell surface (**Figure 7c**) as compared to that of other two glycoproteins (EGFR and DC-SIGN).

**Figure 7:**
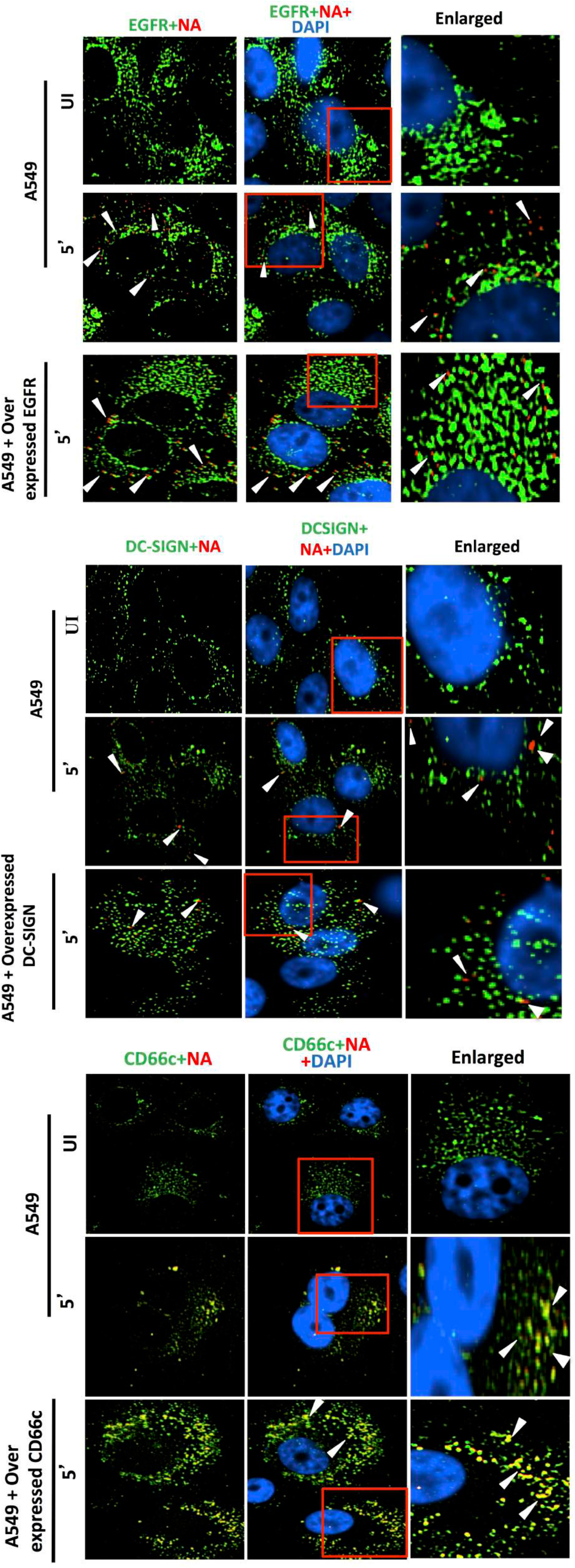
**Comparing ability of host membrane proteins CD66c, EGFR and DC-SIGN respectively in IAV binding on A549 cells through fluorescent microscopy. (a)** Here, UI denotes uninfected cells; 5’, Cells after 5 minutes of virus binding to them. A549 cells were cultured either with endogenous level of EGFR or with overexpression of EGFR (through transfection of EGFR plasmids). Cells were then either left uninfected or infected with 5 MOI of A/PR8/34 influenza virus for 5 minutes. Cells were fixed with 4% formaldehyde for 20 minutes and stained extracellularly (without permeabilization) with anti-EGFR and anti influenza NA antibodies. The secondary antibody Alexa 488 (green) probed host EGFR and Alexa 594 (red) probed influenza NA, cell nuclei is stained with DAPI (blue). The figure shows there is no trace of virus binding in uninfected cells (top panel). However, modest virus binding on cell surface is observed after 5 minutes of incubation as determined by staining influenza neuraminidase NA (red). Interestingly, figure shows that only few virus particles are colocalizing with EGFR (middle panel, yellow spots pointed with white arrow). When EGFR is overexpressed in A549 cells there is neither corresponding increase in virus binding nor in colocalization of viral NA with EGFR. Rather cells overexpressing EGFR shows similar virus binding pattern as cells with endogenous level of EGFR (bottom panel). Altogether, these results demonstrate that virus binding is not significantly increased with overexpression of EGFR, suggesting its poor binding ability with virus. (b) The figure shows A549 cells stained for host DC-SIGN (green) and influenza NA (red) after 5 minutes of virus binding on them. The upper panel of the figure shows no trace of virus binding in uninfected cells. The figure shows few co-localization spots (yellow spots, middle panel) in cells with endogenous level of DC-SIGN. Also, in lung A549 cells overexpressing DC-SIGN (green) there is no consequent increase in virus binding (bottom panel, pointed with white arrow) as compared to cells with endogenous level of DC-SIGN (middle panel, pointed with white arrow). Altogether, results demonstrate that virus binding is not significantly increased with overexpression of DC-SIGN, implying its poor virus binding ability. (c) A monolayer of A549 cells when stained for endogenous expression level of host CD66c (green) and influenza NA (red) after 5 minutes (5’) of virus binding on them, showed significant virus binding on cell surface as probed by influenza neuraminidase NA (red) (middle panel). Cells also showed colocalisation (yellow spots) between endogenous levels of receptor CD66c (green) with influenza neuraminidase NA (red). Unbound influenza NA (red) is not easily identified as most of them are seen merged (as yellow) with CD66c. The upper panel of the figure shows no trace of virus binding in uninfected cells (UI). Also, in A549 cells overexpressing receptor CD66c, a significant increase in virus binding and co-localization of CD66c with NA (yellow dots) on the cell surface is observed (bottom panel) when compared to cells with endogenous level of CD66c (middle panel). The figure shows receptor CD66c (green) mainly at the periphery of cells interacting with significant number of NA (yellow after merge, pointed with white arrow). Altogether, results exhibit that virus binding with CD66c is prominent and significantly increased with overexpression, highlighting the strong binding ability of receptor CD66c as compared to EGFR and DC-SIGN.

An increase in virus binding to receptor leads to consequent virus entry into cells. Therefore, after virus binding experiments, we tested and compared ability of other two glycoproteins (EGFR and DC-SIGN) in virus entry with that of CD66c, at the same experimental conditions. We observed, CD66c overexpression in lung cells resulted significant increase in virus entry as monitored by expression levels of viral NP inside cells (**Figure 8a**). In contrast, overexpression of DC-SIGN and EGFR did not show much change in virus entry, except for a modest increase in viral NP (**Figure 8b, 8c**). We also found that transient overexpression of CD66c did not affect the expression levels of glycoproteins EGFR and DC-SIGN (**Figure 8a**). Similarly, the overexpression of these two glycoproteins (DC-SIGN and EGFR) had no effect on CD66c expression (**Figure 8b, 8c**).

**Figure 8:**
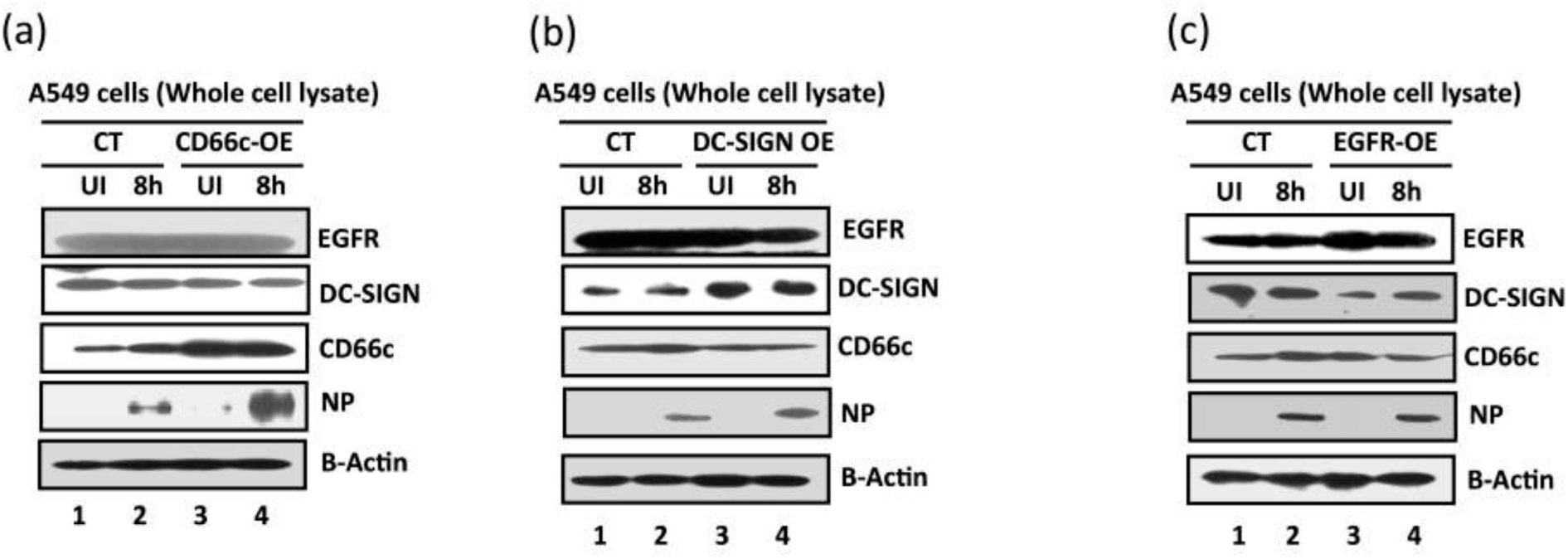
**Comparative measurement of viral infection in cells overexpressing CD66c, EGFR and DC-SIGN respectively.** Here CT denotes control A549 cells (untransfected) with endogenous levels of protein expression; CD66c-OE is A549 cells overexpressing CD66c; DC-SIGN-OE, A549 cells overexpressing DC-SIGN; EGFR-OE, A549 cells overexpressing EGFR; UI, uninfected cell groups; 8h, cells harvested after 8 hours of A/PR8/34 influenza virus infection. **(a)** The A549 cells overexpressing CD66c (CD66c-OE) demonstrate significantly increased virus entry as against A549 cells with endogenous levels of CD66c (CT) when determined by the expression levels of viral protein NP (fourth panel from top). Also, top two panels in figure show that overexpression of CD66c has not affected expression levels of EGFR and DC-SIGN, and their expression levels remained same as in untransfected A549 cells (top two panels). β-Actin is loading control. **(b):** cells overexpressing DC-SIGN (DC-SIGN-OE) showed a slight increase in virus entry into cells as against A549 cells with endogenous levels of DC-SIGN (CT), when determined by the level of viral NP protein (fourth panel from top). More importantly, overexpression of DC-SIGN has not increased expression levels of EGFR and CD66c, whose levels remained same as in untransfected A549 cells (top and third panel from top). β-Actin is loading control here. **(c):** The expression levels of viral NP protein (fourth panel from top) suggests that overexpression of EGFR in A549 cells (EGRF-OE) does not result in further increase in virus entry. The virus entry in these cells is same as in A549 cells with endogenous levels of EGFR (CT). Accordingly, overexpression of EGFR does not increase expression levels of DC-SIGN and CD66c (second and third panel from top). β-Actin is loading control.

### Absence of CD66c in cells showed no decrease in entry of non-IAV virus

The membrane glycoprotein DC-SIGN has been documented to serve as a low-specificity virus receptor for IAV and is postulated to facilitate the entry of other viruses like HIV (Human immunodeficiency virus) and HCV (Hepatitis C virus) (11). Critically, we argued that CD66c being a membrane glycoprotein might also be expected to serve as a low-specificity receptor for viruses other than IAV. To this effect, we sought to establish the specificity of CD66c towards influenza virus against an unrelated RNA virus – HCV. We conducted these experiments in human hepatoma Huh cells that were siRNA-mediated-silenced for CD66c expression. We monitored HCV entry into these Huh cells by checking the expression levels of the HCV NS3 protein (**Figure 9**). This data clearly showed that the absence of CD66c in Huh cells had not inhibited entry of HCV, thus proving that CD66c was not a low-specificity general viral receptor.

**Figure 9:**
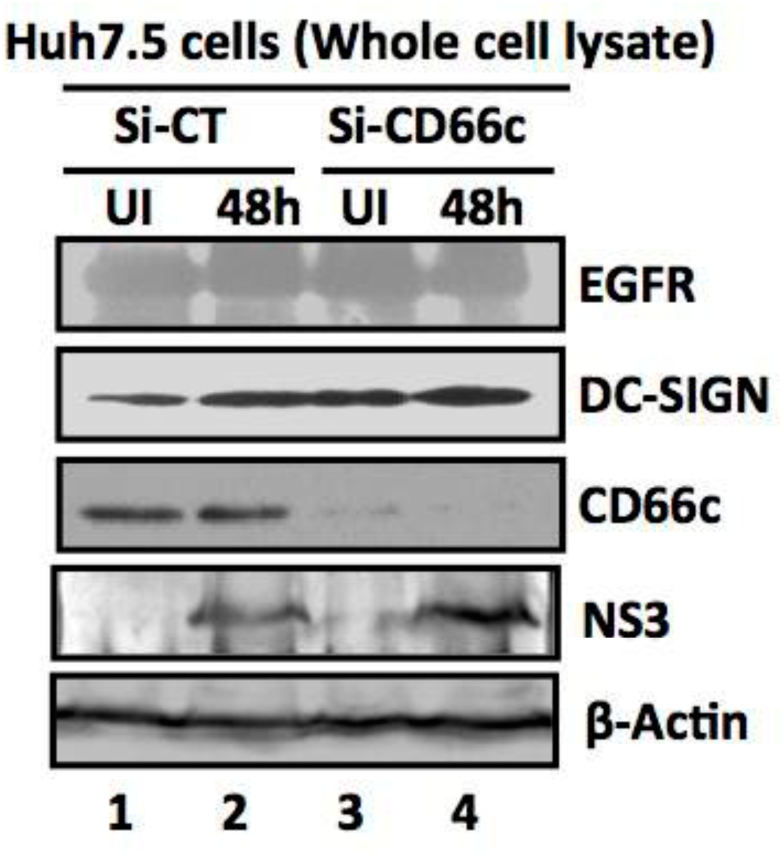
**siRNA knockdown of CD66c in Huh7.5 cells have not inhibited entry of another non-IAV virus (Hepatitis C virus, HCV).** Here, Si-CT denotes control siRNA treated human hepatoma cells; Si-CD66c is human hepatoma cells treated with CD66c siRNA. Cells are either left uninfected (UI) or infected with 0.5 MOI of HCV for 48 hours (48h). Human hepatoma cells harvested 48 hours post infection is subjected to immunoblotting. The figure shows that CD66c siRNA treated human hepatoma Huh7.5 cells show reduced expression of CD66c (third panel from top). However, this siRNA-mediated knockdown of CD66c in Huh7.5 cells does not inhibit HCV entry as determined by the expression level of HCV viral protein NS3 (fourth panel from top). Also, Huh7.5 cells knocked down for CD66c expression does not show any inhibitory effect in the expression levels of EGFR and DC-SIGN (upper two panels). β-Actin is loading control.

### Influenza virus incubated with CD66c showed reduction in alveolitis

Since we had clearly shown that CD66c was capable of binding to the NA of IAV, we reasoned if we could use heterologously expressed recombinant CD66c protein to bind IAV particles, this should in principle bring down the infectivity of the virus in mice. Thus, we incubated 1μg of biologically active recombinant CD66c (rCD66c), that was produced in mouse myeloma cell lines, with 7.4 × 10^7^ PFU IAV before intranasal infection of BALB/c mice. After ten days of infecting mice with virus through intranasal inoculation, we noticed a considerable reduction in alveolitis in the mice that were infected with rCD66c bound IAV (**Figure S1a**) as compared to mice treated with virus alone (**Figure S1b**) or virus incubated with protein control Bovine Serum Albumin (BSA) (**Figure S1c**). The insignificant inflammation in the lungs of mice infected with CD66c treated virus was comparable to the lung tissues from uninfected mice (**Figure S1d**). These results strongly suggest that binding of rCD66c to NA significantly reduces lung pathology of IAV infected BALB/c mice, thus confirming our belief that CD66c is a receptor for IAV. The incubation of 1μg of biologically active recombinant CD66c (rCD66c) with 7.4 × 10^7^ PFU IAV had shown inhibition of virus entry in human A549 lung cell line (Data not shown).

## Discussion

In our search for a receptor for IAV, we chose to follow the experimental path followed by many other research groups to identify new viral receptors (13-20). We conducted similar experiments in detail, which could validate the interaction between viral NA and host CD66c at the outer cell surface, during IAV attachment and entry. The results thus obtained from these experiments provided sufficient evidence that suggested CD66c as the glycoprotein receptor for influenza virus. The validation of a protein receptor for influenza virus from this study provides valuable insights into some unresolved problems of influenza entry. For instance, it was cited in the earlier studies on influenza entry that although sialic acid was required for virus binding, a specific subset of glycoprotein receptors was necessary for effective viral entry (8, 9), which is yet unknown. Therefore, with the results presented here, we suggest that CD66c is at least one, of the possibly many, glycoprotein receptors. Additionally, this study may further lead to the discovery of other glycoprotein receptors or co-receptors playing a role in virus entry besides CD66c. The mechanism of virus entry is poorly understood and different alternative routes were suggested for IAV entry, such as clathrin-mediated endocytosis, non clathrin-mediated, caveolin-mediated endocytosis or macropinocytosis (21-28). We believe, CD66c, as a receptor for the influenza virus will help in elucidating the precise route for virus-entry and pave the way for discovery of other co-receptors and their mechanism.

Further, with this finding we noticed that IAV follows an infection pattern that is similar to some other viruses wherein they take advantage of adhesive properties of hosts cell adhesion molecules (CAMs), for their attachment and entry. For example, Coronavirus, Rabies, Reovirus and Rhinovirus, employ the following cell adhesion molecules CEACAM1, NCAM-1, JAM-A, ICAM-1 respectively, as their receptor for cellular entry (13-15). Additionally, viruses often interact and utilize these cell adhesion molecules (CAMs) to foster a contact between infected and uninfected target cells for an effective cell-to-cell spread (29). Carcinoembryonic Cell Adhesion Molecule 6 (CEACAM6/CD66c) as the receptor for IAV opens opportunities for further investigations on cell-to-cell spread for this virus as well. Our result showing prominent NA-CD66c interaction at the site of cell-cell junction compared to the rest of the cellular membrane is a preliminary indication in that direction (**Figure 1c**).

Apart from cell adhesion other unique attributes of CD66c, such as GPI anchoring, lipid raft association and heavy glycosylation (Sialyl-Lewis^X^), make it a very suitable and strong receptor candidate for IAV entry. Influenza binding on the cell surface causes lipid raft mediated virus uptake (*10*), hence we suggest that this putative receptor CD66c being a component of lipid rafts bears potential to further dissect and solve the enigma of the viral internalization mechanism. In addition to that, Sialyl-Lewis^X^ is reported as the common receptor determinant of a number of influenza viruses of the terrestrial poultry (*30*). Therefore presence of Sialyl-Lewis^X^ on the CD66c molecules makes the latter a strong glycoprotein receptor candidate for the IAV. More importantly, in human lungs there is abundant expression of CEACAM6 by alveolar and bronchial epithelial cells, where it also demonstrates surfactant association and secretion into lung-lining fluid (31). These features of CD66c with respect to human lungs make this molecule vulnerable to respiratory pathogens like IAV. More importantly, like other CEACAMs, which are receptors for respiratory pathogens (bacteria) including *Haemophilus influenzae* and *Moraxella catarrhalis* (32, 33), CEACAM6 (CD66c) from above results, serves as a receptor for yet another respiratory pathogen - IAV.

It is reported that when pathogens interact with the CEACAM receptors, there is significant activation of PI3K signaling during internalization of the pathogen (34). In our previous report, we validated the activation of PI3k/Akt pathways when CEACAM6 (CD66c) interacts with influenza NA (12), here we demonstrate viral internalization upon NA-CD66c interaction at the cell surface. Also, this new finding on CD66c provides support to the viewpoint of a contentious argument made in the past on a role for NA in influenza entry (*35, 36*).

More importantly, viruses frequently exploit chemokine receptors, some CD markers and other membrane glycoproteins of IgSF as their receptors for entry and also for manipulating the host defense mechanism (37). For this reason, a majority of these immunomodulatory studies get direct reference to viral infections or immune evasion, at the entry stage, which are to a great extent, centric to the interaction of the viral spike glycoproteins with such cellular receptors and co-receptors. It is important to mention here that, the receptor CD66c is also a member of IgSF and plays a number of crucial immunomodulatory roles in human. To cite a few examples, during multiple myeloma CD66c inhibits cytotoxic T cell activation, in normal neutrophils it is known as an activation marker that stimulates neutrophil signaling (38, 39). Further, CD66c increases apoptosis in B-cell precursor acute lymphoblastic leukemia cells (40). Another set of evidence on the expression of CEACAMs in human lung and their modulated co-expression by type I and type II interferon was reported recently (33, 41). Altogether these studies establish CD66c as an active immunomodulatory molecule playing a significant role in innate and adaptive immunity in human. Accordingly, for being an active immunomodulatory molecule and with presence on T cells and Macrophages, we took elements of caution into consideration while designing animal experiments and interpreting data thereof (**Figure S1**).

Therefore we argue that immunomodulatory studies carried out earlier in abeyance of any glycoprotein receptor for IAV were rather incomplete and CD66c at the helm of immunomodulation and its interaction with virus during internalization has potential to unfold the precise mechanism of influenza infection, consequent immune response and cell tropism. Lack of any identified glycoprotein receptor during IAV attachment and entry, had greatly limited influenza research discourse in this direction whereas similar questions had been addressed well with other viruses (42).

## Material and methods

### Plasmid constructs, antibodies, virus strains, and mammalian cell lines

pRc/CMV plasmid with full-length untagged CD66c gene was used for expression in mammalian cell lines. Human CLEC4M/DC-SIGNR/CD299 gene cDNA ORF clone (cat. HC00654) was purchased from ACROBiosystem Co. LTD. EGFR-GFP plasmid (in EGFP clontech vector backbone) was gifted from Professor Maddy Parson, King’s College London. Monoclonal anti-CD66c antibody (mAb anti-CD66c) was purchased from Santa Cruz Biotechnology, Santa Cruz, CA (Catalog # sc-59899) and anti-NA (αNA) antibody was purchased from Meridian Life Sciences (Saco, ME). Secondary antibody anti-mouse Alexa Fluor^®^ 594 was used against mAb anti-CD66c and anti-rabbit Alexa Fluor^®^ 488 against αNA. Anti-CD209/CD299 (DC-SIGN/L-SIGN) was purchased from BioLegend. A/Puerto Rico/8/34 (PR8) influenza virus strain was used for virus infection experiments both in mammalian cell lines and mice. For detection of viral M1 protein, in-house raised antisera against M1 VLP was used (*43*). For detection of NP in flow cytometry and confocal experiments FITC conjugated anti-NP from abcam^®^ was used (Catalog # ab20921). Influenza virus NP protein was detected using rabbit antisera raised against purified and disrupted PR8 virus (44), which was a kind gift from Dr. Balaji Manicassamy (UIC, Chicago). Virus was used at multiplicity of infection (m.o.i.) of 1 unless where specified. Cell lines such as, Human lung adenocarcinoma epithelial (A549) and NIH3T3 mouse embryonic fibroblast, were purchased from ATCC. Lec2CHO cell line was gifted by Dr. S. Gopalan Sampath Kumar, National Institute of Immunology, New Delhi. Experiments related to human hepatoma cell line, Huh7.5 and JFH-1 infectious HCV were conducted in Dr. Waris’s lab. For biochemical experiment in BALB/c mice we used purified recombinant CD66c, expressed in mouse myeloma cell line, purchased from R&D systems (catalog # 3934-CM, activity checked by the manufacturer).

### RNA interference and virus infection experiments

Mock-infected and PR8-infected A549 and Huh 7.5 cells were transfected with CD66c siRNA according to the manufacturer’s protocols (Santa Cruz Biotechnology). A concentration of 30nM of siRNA was used. The Huh 7.5 cells treated with Si-RNA were cultured into 6 well plates for 48 h then either left uninfected or infected with 0.5 MOI of HCV in the incomplete Dulbecco’s Modified Eagle’s Medium (DMEM) medium for 5 hours, then replaced with complete DMEM medium, followed by incubation for 48 hours in the 5% CO2 incubator at 37°C. Similarly, A549 cells were infected with PR8 virus but with different MOI (1-5), MOI 1 for virus infection level detection and MOI 5 for the virus binding experiments (also specified in legends to figures).

### Antibody mediated receptor blockade experiments and influenza infection to the cells

Cells (A549 and NIH3T3) were plated at a density of 10^6^/well in a 6-well culture plate. The cell monolayers were washed with PBS three times and incubated either with anti-CD66c mAb or with IgG1 isotype antibody, in 200μL of PBS with 3% fetal calf serum (FCS) for 40 minutes at 20°C. Unbound antibody was removed by washing cells with PBS following which cell monolayer was incubated with PR8 virus in OPTI-MEM (3% FCS) for 1 h at 37°C under 5% CO_2_. The medium was replaced with DMEM (10% FCS) and cells were incubated at 37°C under 5% CO2. Cells were harvested for time points of 8 h.p.i or 24 h.p.i. The extent of viral infection in cells was determined by probing viral proteins (M1 and NP) through flow cytometry and western blot analysis.

### Immunofluorescence microscopy

Intracellular and extracellular immuno-staining of cells were performed during this study. For intracellular staining, cell monolayer, that was fixed by overlaying 4% paraformaldehyde, was permeabilised by treating with PBS containing 0.5% Triton X-100 for 10 minutes at 37°C. Following this, the cell monolayer was conditioned and blocked with Phosphate-buffered saline (PBS) containing 0.5% Triton X-100 and 0.5% [wt/vol] Bovine serum albumin (BSA) for 1 h. Permeabilised cells were incubated with FITC-conjugated primary antibody against viral NP at a dilution of 1:200 in antibody solution (PBS containing 0.5% Triton X-100, 0.1% [wt/vol] sodium azide and 0.5% [wt/vol] BSA) for 30 min. This was followed by washing cells with PBS containing 0.5% Triton X-100 to remove excess antibodies and mounting cells on slides for observation under confocal microscope (A1R; Nikon, Tokyo, Japan). For extracellular staining was two-step process, wherein cells fixed in 4% paraformaldehyde were incubated with primary antibody (anti-CD66c or anti-NA) followed by respective fluorescent secondary antibodies, bypassing the cell permeabilisation step. Primary antibody was used at a dilution of 1:100 and secondary antibodies were used at a dilution ratio of 1:1000 in antibody solution.

### Flow cytometric analysis

For intracellular Influenza NP staining, viral infected cells were washed with PBS and centrifuged to remove debris. Single cell suspension thus formed was fixed with 4% paraformaldehyde (PFA) in PBS. Fixed cells were then washed twice with PBS containing 3% FCS and resuspended in a permeabilisation buffer (Cytofix/Cytoperm kit; BD) for 10 minutes. The permeabilised cells were incubated for 45 minutes with Fluorescein isothiocyanate (FITC) conjugated NP antibody (primary) in PBS (with 3% FCS). The stained cells were washed with buffer (3% FCS in PBS) to remove unbound antibody and taken for cytometric readings. For extracellular staining of CD66c or the viral NA attached to cells during virus binding experiments, we performed standard extracellular staining protocol bypassing the cell permeabilisation step. Cells with or without virus on their surface were fixed with 4% PFA in PBS, followed by washing with PBS. Fixed cells or cell-virus complex were then incubated with anti-CD66c primary monoclonal antibody or anti-NA primary antibody in 50 μL of PBS with 3% FCS for 40 minutes at 4°C (antibodies to solution ratio was 1:200). After washing unbound antibodies cells were incubated with secondary antibody against these primary antibodies (anti-CD66c, anti-NA) in 50μL of PBS with 3% FCS for 40 minutes at 4°C. Ratio of secondary antibodies to solution was 1:1000 (v/v). After washing unbound secondary antibodies to cells in PBS with 3% FCS, stained cells were taken for cytometric analysis. Fluorescence intensity was measured by flow cytometry (FACS Calibur; BD) and data was analysed using FlowJo (Tree star, USA).

### Semi quantitative and real-time pcr

Total RNA from cells was extracted using RNeasy Mini Kit (Qiagen) and treated with DNase I (Invitrogen). 2μg of RNA was reverse-transcribed using M-MLV Reverse Transcriptase (ThermoFisher, Catalog # 28025013) in a volume of 20 μL. The synthesized cDNA was diluted 1:5 in water. 2.0 μL of cDNA was then used in a SYBR^®^ Green PCR Master Mix (Applied Biosystems) based real-time PCR reactions in a volume of 20 μL. StepOne^TM^ PCR machine was used to acquire real-time PCR readouts.

Primer sequence: Following primer sequences were used for the semi quantitative and Real-time pcr reactions.

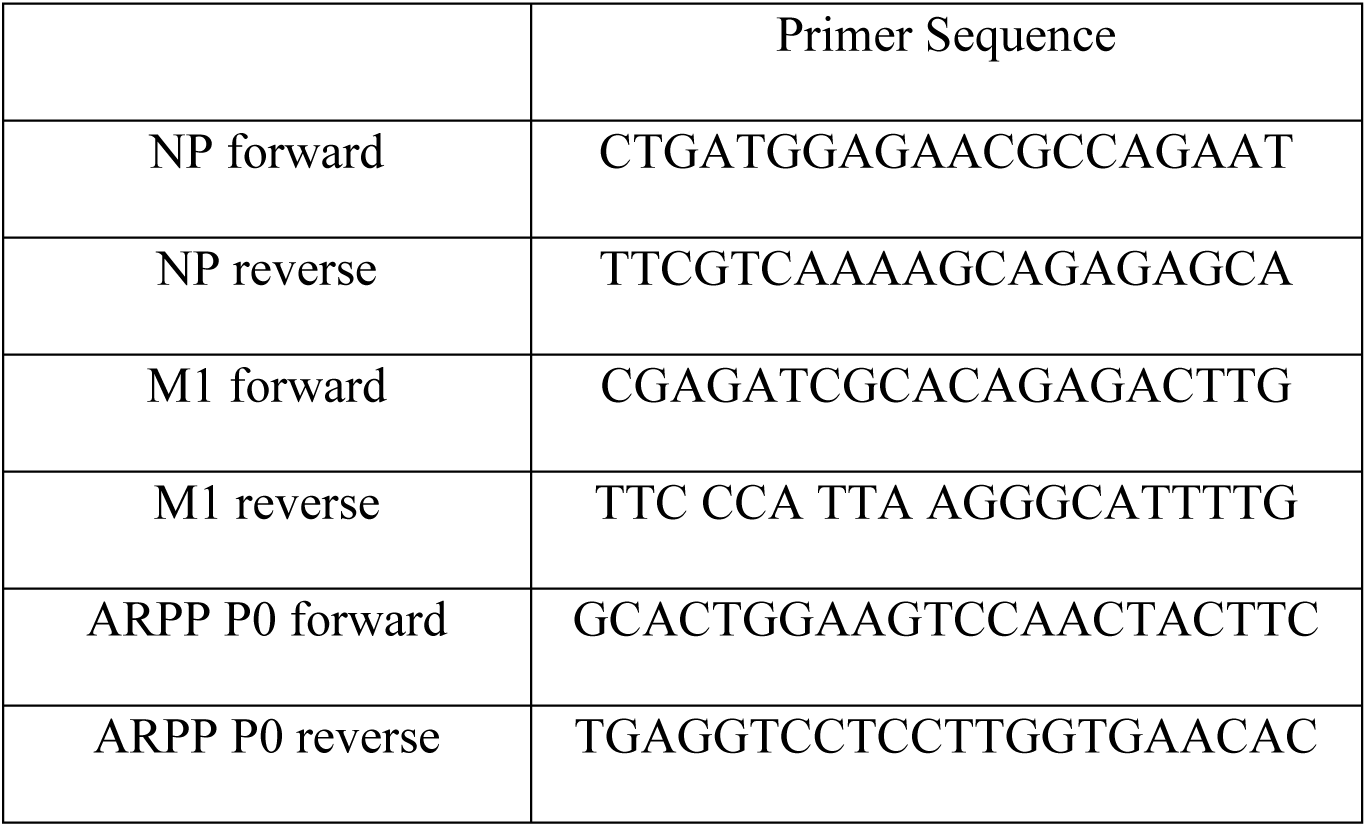

### Incubation of virus with recombinant protein, intranasal challenge of mice and lung histology

1 μg of recombinant CD66c protein was incubated with 50 μL of PR8 virus in PBS, with a titer of 1.48 × 10^9^ pfu/mL, for 30 minutes so that protein binds with virus. Six-week-old female BALB/c mice were first anaesthetized with isoflurane, and were inoculated with 50 μL of virus (incubated with recombinant CD66c), in PBS intranasally. Mock mice group were inoculated with 50 mL of virus pre-incubated with BSA as a protein control. Also a group of mice was inoculated with untreated PR8 virus (virus without any protein incubation). Survival and the body weight of the mice were monitored regularly for 10 days post infection. Mice were then euthanized and lungs were taken out for study. Lung tissues were preserved in formalin embedded in paraffin and were cut in uniform 4-μm sections. Tissue sections were stained with hematoxylin and eosin stain and examined for histopathological changes under the microscope at 20X magnification.

## The abbreviations used are

SIA, sialic acid; IAV, influenza A virus; EGFR, epidermal growth factor receptor; L-SIGN, Liver/lymph node-specific intercellular adhesion molecule-3-grabbing integrin; DC-SIGN, Dendritic cell-specific intercellular adhesion molecule-3-grabbing nonintegrin; GPI-anchored, Glycosylphosphatidylinositol-anchored; HA, Hemagglutinin; NA, neuraminidase; PI3K, phosphatidylinositide 3-kinases; NA-CD66c, interaction between Neuraminidase and CD66c; NP, nucleoprotein of IAV, M1, a matrix of IAV; CAMs, cell adhesion molecules; IgSF, immunoglobulin superfamily; mAb, monoclonal antibody; CEACAM6, carcinoembryonic antigen-related cell adhesion molecule 6; CD66c, cluster of differentiation 66c; m.o.i., multiplicity of infection; h.p.i, hours post infection; DAPI, 4’, 6-diamidino-2-phenylindole; HCV, Hepatitis C virus.

## Authors’ contributions

S.K.R. and S.K.L. conceived, planned and orchestrated the project. S.K.R. planned and optimized antibody mediated cell blockade experiments in human lung A549 cells. S.K.R performed other experiments carried out in A549 and lec2CHO cell lines. P.G. performed experiments in NIH3T3 cell lines. M.A.A. performed comparative binding, entry of IAV with reference to CD66c, EGFR, DC-SIGN and siRNA (against CD66c expression) experiments to compare entry of IAV and HCV. I.A., C.C., S.S., and S.C performed virus experiments in mice. D.K.V. performed virus entry experiment in HEK cells. S.K.R. prepared first draft of the manuscript. S.K.R., N.N and S.K.L further improved the manuscript from inputs of other authors. S.K.R., D.W., G.W., and S.K.L critically analyzed the datasets.

## ACKNOWLEDGEMENT

S.K.R was awarded CSIR JRF funding. The plasmid pRc/CMV with untagged CD66c was a generous gift from Wolfgang Zimmermann (Tumor Immunology Laboratory Life Center, University Clinic-Grosshadern Muenchen, Germany). The authors thank Professor Maddy Parsons, King’s College London for EGFR-GFP Plasmid. We thank Dr. Aftab Ahmad, University of Alabama at Birmingham for A549 cells, Dr. Joseph Reynolds for PR8 whole virus and Balaji Manicassamy (UIC, Chicago) for antisera raised against purified and disrupted PR8 virus. The authors thank Purnima Kumar (ICGEB), Ravinder Kumar (ICGEB) and Jyoti Batra (ICGEB) for their help with the reagents and media preparation. Initial support from Sultan Tousif (ICGEB) during flow cytometry measurements is duly acknowledged.

---

**Figure S1:**
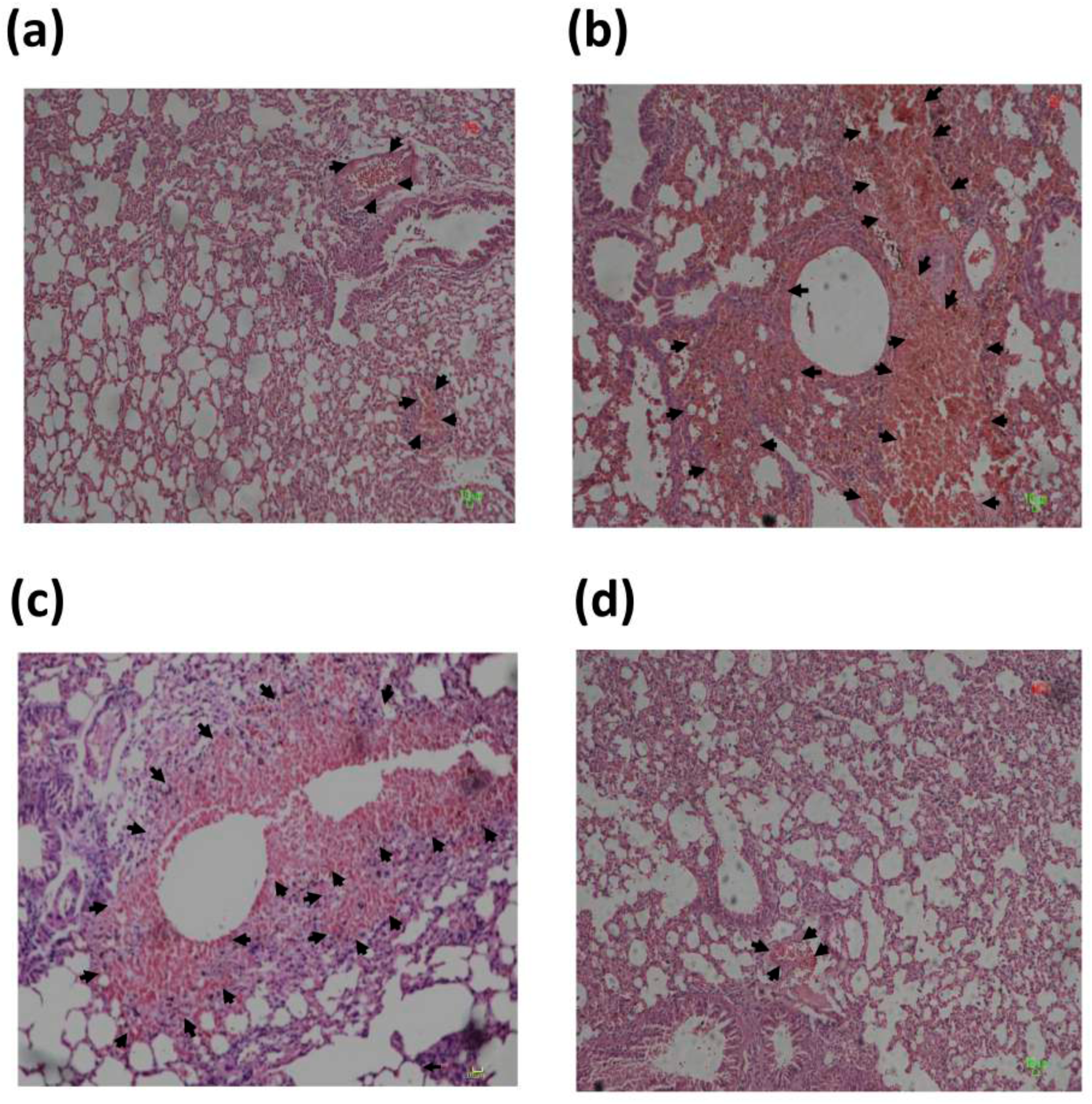
**PR8 virus pre incubated with purified recombinant CD66c causes lower inflammatory response in mice lung.** Figure shows histopathological analysis of lung tissues of mice infected with A/PR8/34 influenza virus. Images shown in the figure are lung tissues of sacrificed animals, preserved in formalin, embedded in paraffin and sectioned into serial 4-µm sections. The figure shows images of infected tissues stained with hematoxylin & eosin dye (H&E) captured at 20X magnification. The representative images of mice from different experimental animal groups are — **(a)** the lung tissues from mice infected with A/PR8/34 virus pre-incubated with CD66c that showed a mild alveolitis and slight hemorrhage (indicated with arrow). **(b)** H&E stained lung tissues from mice infected with A/PR8/34 virus, showing significant alveolitis and extensive intra-alveolar hemorrhage (marked by arrow). **(c)** H&E stained lung tissues from mice infected with A/PR8/34 virus pre-incubated with BSA (as a mock protein control), showing alveolitis and intra-alveolar hemorrhage (marked by arrow). **(d)** Lung tissues from uninfected mice showing no sign of alveolitis.

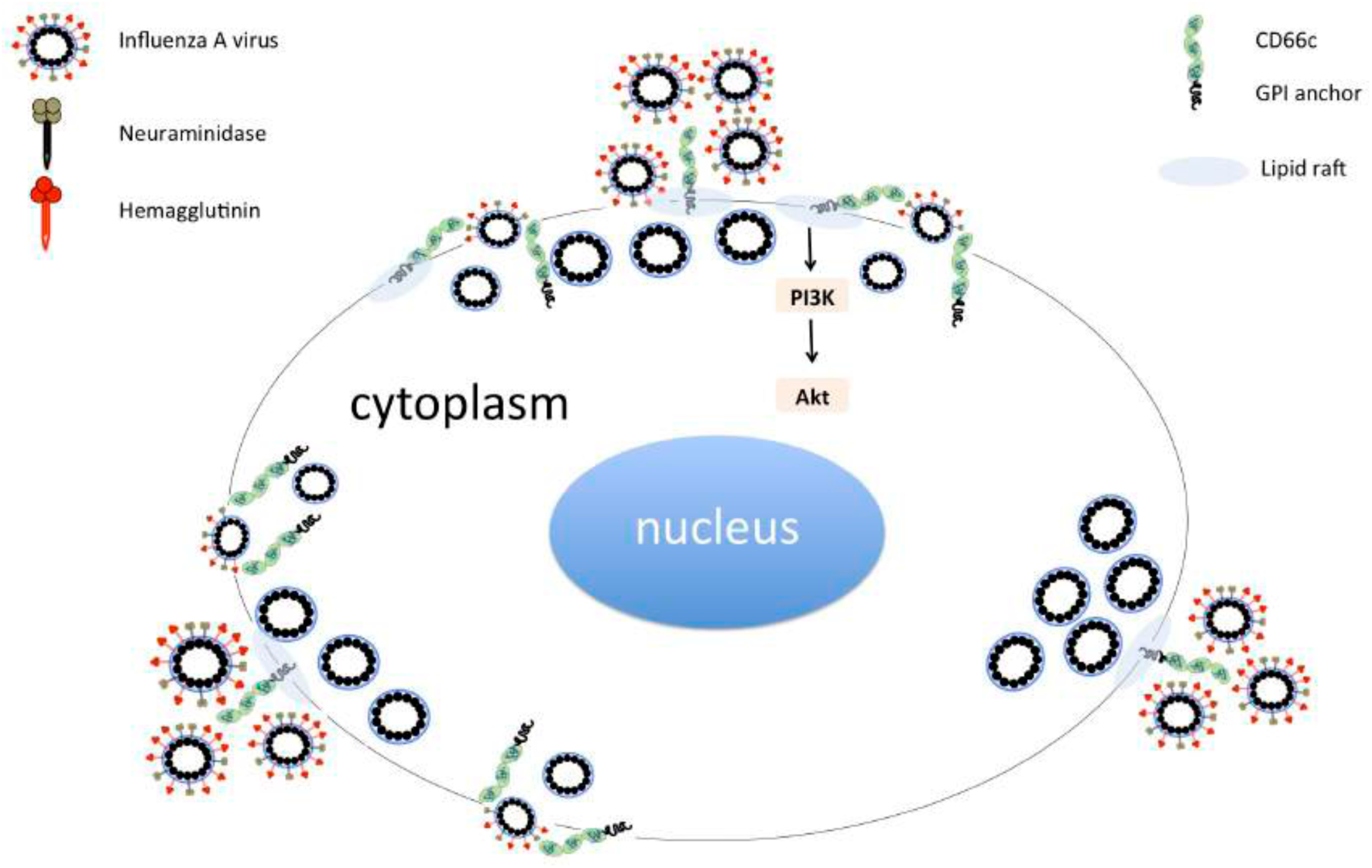
Graphical abstract.

